# Identify phage hosts from metaviromic short reads based on deep learning and Markov chain model

**DOI:** 10.1101/2021.03.01.433351

**Authors:** Jie Tan, Zhencheng Fang, Shufang Wu, Qian Guo, Xiaoqing Jiang, Huaiqiu Zhu

## Abstract

Phages - viruses that infect bacteria and archaea - are dominant in the virosphere and play an important role in the microbial community. It is very important to identify the host of a given phage fragment from metavriome data for understanding the ecological impact of phage in a microbial community. State-of-the-art tools for host identification only present reliable results on long sequences within a narrow candidate host range, while there are a large number of short fragments in real metagenomic data and the taxonomic composition of a microbial community is often complicated. Here, we present a method, named HoPhage, to identify the host of a given phage fragment from metavirome data at the genus level. HoPhage integrates two modules using the deep learning algorithms and the Markov chain model, respectively. By testing on both the artificial benchmark dataset of phage contigs and the real virome data, HoPhage demonstrates a satisfactory performance on short fragments within a wide candidate host range at every taxonomic level. HoPhage is freely available at http://cqb.pku.edu.cn/ZhuLab/HoPhage/.

## 1. Introduction

Viruses are the most abundant organism on earth and phages - viruses that infect bacteria and archaea - are dominant in the virosphere (Breitbart *et al*., 2005) and play an important role in the microbial community (Shkoporov *et al*., 2019). To explore the ecological impact of phage in a community, it is imperative to assign the host of a given phage (de Jonge *et al*., 2019). With the help of metagenomics technology, a wealth of novel phages that cannot be cultured are identified. Compared to the traditional culturing-based approach which naturally carries direct host information, the metagenomic method, especially metavirome, lacks the links between phages and their hosts, thus brings the increasing demand to develop computational tools for host identification of short phage fragments. However, the relevant research is still insufficient until now.

Recently, several computational strategies, mainly based on abundance profiles, genetic homology, CRISPRs, exact matches, and oligonucleotide profiles, have been proposed for host identification (Edwards *et al*., 2016). Some of these strategies rely on a known database. For example, since some phages can integrate their genomes into host chromosomes, a prokaryote containing homologous regions with the given phage may be the potential host. Also, some prokaryotes can incorporate some DNA fragments from phages that have infected them into their own genome forming interspaced short palindromic repeats (CRISPR) spacers, hence searching the CRISPR spacers on a bacterial genome can help to identify the phage which can infect it if it contains the CRISPR system. Another strategy is based on the abundance profiles. Because phages cannot thrive without their host, the bacterium which has a good correlation in abundance with a phage across multiple samples may be the host of it. However, such approaches mentioned above present poor performance in metagenomic data. The microbial community contains a large number of novel phages that have low similarity with the known phages in the current database (Hayes *et al*., 2017), and therefore, a similarity search-based approach cannot handle the task of host prediction of novel phages. Also, a strategy based on CRISPR spacers only works well when the phage fragment is long enough to cover the CRISPR region, hence it is not suitable for metagenomic data which contains a large number of short fragments. Moreover, a strategy based on abundance profiles requires multiple samples to calculate the correlation between each phage and bacterium, and the population dynamics between phages and their hosts often present non-linear behavior (Edwards *et al*., 2016), increasing the difficulty of host identification.

In contrast, strategies based on sequence signatures are more suitable for metagenomic data. Phages must survive together with their host and are under strong selection pressures during the phage/host co-evolution. As the result, phages will adapt the sequence signatures of their host, such as GC content and codon usage, to its host (Edwards *et al*., 2016). It is considered that sequence signatures between phage and host or between phages that infect the same host are similar. Since prokaryotes of different genera often contain different sequence signatures, it is thus considered that sequence signatures can be used to classify the infection relationship between a phage sequence and a specific prokaryote. For instance, a research has shown that k-mer frequencies can be used to identify some phages whose hosts belonging to certain genera (Zhang *et al*., 2017). Because sequence signatures often distribute over the phage whole genome, identifying the host of a given phage fragment using sequence signatures is relatively effective. Also, such an approach does not rely on sequence alignment so it can identify hosts of fragments that derive from novel phages. Currently, some computational tools based on sequence signatures have been developed. VirHostMatcher (Ahlgren *et al*., 2017) calculates the oligonucleotide frequency dissimilarity between a phage sequence and each candidate host and the one with the lowest dissimilarity will be selected as the predicted host. WIsH (Galiez *et al*., 2017) uses the Markov chain model to calculate the similarity between phage sequence and each candidate host, and similar to VirHostMatcher, the one with the highest similarity is selected as the predicted host. VirHostMatcher-Net (Wang *et al*., 2020) is an upgraded tool that integrates multiple features, including CRISPR spacers and alignment-free similarity measures used in VirHostMatcher and WIsH. However, the performance of these tools in short DNA fragments generated by large-scale sequencing technology is rather unsatisfactory. Moreover, with the expansion of candidate host range, the accuracy of these tools decreases drastically. As the state-of-the-art tool for short phage fragments, WIsH only acquires an accuracy at the genus level of about 60% for 3,000 bp fragments among 20 candidate host genera while a considerable proportion of assembled contigs in the metagenomic data obtained by next-generation sequencing are shorter than 3,000 bp (Smits *et al*., 2014).

Machine learning algorithms, especially deep learning algorithms, have been widely used to infer the relationship between two biological elements in the field of bioinformatics, such as gene-gene relationships (Yuan *et al*., 2019), protein-protein interaction (Hashemifar *et al*., 2018), RNA-protein interaction (Yang *et al*., 2018). Deep learning algorithms have been further applied to the host prediction of a virus. VIDHOPHAGE (Mock *et al*., 2020) is a deep learning-based virus-host prediction tool that obtains highly accurate predictions on three different virus species while using only fractions of the viral genome sequences. Markov chain model is also a popular method for researches of biological sequence. WIsH (Galiez *et al*., 2017) is a phage host prediction tool that uses the Markov chain model, and obtains relatively good performance on short phage fragments. The availability of the Markov chain model used by WIsH is also validated by VirHostMatcher-Net (Wang *et al*., 2020) since it integrates multiple features, including the score of WIsH, to predict the host of short phage fragments.

Considering the phage fragment in the metagenomic data of real community is short in length, as well as the taxonomic composition of microbial community is complex, we developed HoPhage (Host of Phage), a tool of host identification for a given phage fragment, which demonstrates high performance on short fragments within a much wider candidate host range. HoPhage integrates two modules, HoPhage-G and HoPhage-S, to improve performance, G and S respectively mean that the model is built at the genus level and the strain level. HoPhage-G is a deep learning-based module to judge whether the given phage fragment can infect any prokaryote from a specific genus. By constructing pairs of phage fragments and prokaryotes at the genus level, host identification is transformed from a complex multi-class prediction issue to a binary classification task of judging whether there is an infection relationship between a pair. In order to improve model performance, we adopted the inception module from GoogLeNet (Szegedy *et al*., 2016), which can extract features at multiple scales. However, the distribution of the number of phages infecting different host genera is uneven in the current database and a large proportion of known phages derives from a narrow host range (Roux *et al*., 2016). As we all know, machine learning methods rely on the existing data that is used for training. So as a complement, we incorporated HoPhage-S which is a Markov chain model-based module to address this challenge. HoPhage integrates the scores of these two modules by calculating the weighted average score of them and select the host genus with the highest score as the final prediction. Testing on the benchmark dataset of artificial phage contigs and real virome data, HoPhage demonstrates superior performance on short fragments within a wide candidate host range at every taxonomic level. HoPhage is freely available at http://cqb.pku.edu.cn/ZhuLab/HoPhage/ or https://hub.docker.com/repository/docker/jietan95/hophage.

## 2. Material and Methods

### 2.1. Benchmark datasets and training-test splitting

We first downloaded 5,314 complete prokaryote genomes which are included in the KEGG GENOME database from NCBI through its GeneBank accession number. These 5,314 prokaryote genomes range from 1,192 different genera while 100 genomes have no genus annotation. ‘prokaryotes_info.csv’ is the information list of these prokaryote genomes and is available in http://cqb.pku.edu.cn/ZhuLab/HoPhage/data/.

We then retrieved all the 3,106 phages included in Virus-Host DB (Mihara *et al*., 2016) from NCBI and kept phages that can infect the prokaryote belonging to the genera with at least 2 phages are annotated to infect them. At last, 2,965 genomes of phages whose host range from 155 genera are used in our study. The accession number list of these phage genomes and their host annotations are shown in ‘phages_hosts_info.csv’ and the distribution of host genera according to the number of phages annotated to infect them in VirusHost DB is shown in ‘genus_distribution.csv’.

As an ab initio tool, it is important to evaluate whether the algorithm can identify the hosts of the fragments from a novel phage. Due to the lack of phage fragments with detailed host annotations from experimental metagenomics, we used MetaSim (Richter *et al*., 2008) to generate a benchmark dataset of artificial short contigs after the whole genomes of phages were downloaded from the NCBI database. For HoPhage-G, to ensure that all test data is “novel” for the model, we split the training and test set of phage genomes by the date when the phage was firstly published before we simulated artificial fragments. In general, phage genomes released before 2015 were used to train the deep learning model, and those released after 2015 were used to evaluate the algorithm. After this decisive partition, the ratios of the number of genomes in the training set to the test set of some genera were unreasonable compared with the traditional ratio. Therefore, we manually adjusted the genomes of phages that can infect these genera to meet a reasonable proportion of genomes in training and test data. To ensure that the training and the independent test set do not have identical or near-identical phage genomes, we followed these two principles during the adjustment process: 1) the same phage genomes will not exist in the training set and test set at the same time; 2) different phage genomes from the same study must exist in the training set or test set at the same time. For HoPhage-S, all prokaryote genomes in our dataset were used to construct the codon Markov chain model. Since there is no need to use phage genomes to construct a Markov chain model, all test set used in HoPhage-G were kept to evaluate the performance of HoPhage-S.

In the first place, three groups of phage fragments with different length intervals were constructed separately, which were 100-400 bp, 401-800 bp, and 801-1,200 bp. Since the input length of the deep learning model is fixed, we independently constructed training sets of corresponding length intervals to train the deep learning model for each group in HoPhage-G. Besides, test sets of each group were also simulated independently to evaluate the performance of HoPhage. As a result, we constructed 500,000 training fragments and 5,000 test fragments for each group. Subsequently, two test sets of longer phage fragments were constructed to further verify the effectiveness of HoPhage, which are 1,201-3,000 bp, 3,001-5,000 bp.

### 2.2. HoPhage-G: the deep learning model based module

In the existing dataset we used in this study, the number of phages that can infect a specific genus varies greatly, which can be seen from the distribution list in ‘genus_distribution.csv’. For deep learning or other machine learning algorithms, it is difficult to train a multi-classification model using a quite unbalanced dataset, especially when there is no sufficient data for some classes. It has been pointed out that the hosts of most phages have specificity at the genus level (Koskella *et al*., 2013). We counted all 3356 host annotations of phages in VirusHostDB and found that only 72 (2.15%) annotations lack the genus-level information, and 278 (8.28%) annotations lack the species-level information, but 2599 (77.44%) annotations miss strain-level information. Since it is easy to understand that the host prediction of phages can be regarded as the interaction prediction between phages and their candidate hosts, that is, to determine whether there is an infectious relationship between the phage fragment and a potential host. It is more appropriate that the pairs are constructed at the genus level for the above reasons. Therefore, in HoPhage-G, this complex multi-classification host prediction issue transformed into a two-classification task through constructing pairs of phage fragments and genera of prokaryotes. This module was named HoP-G because the pair was constructed at the genus level.

The process of data preparation for HoPhage-G is mainly the transformation of phage fragments and the extraction of sequence features of prokaryote genomes (Fig. 1A). For phage fragments, we first extracted artificial contigs from the phage whole genomes in the training set and the test set respectively by MetaSim. Each artificial contig was represented by “one-hot” encoding form, namely, base A, C, G, T was represented by the “one-hot” vector of [1,0,0,0], [0,1,0,0], [0,0,1,0] and [0,0,0,1]. As mentioned earlier, the length of the phage fragments in the simulated data set we constructed varies within 3 length intervals, and the input length of the deep learning model is fixed. Hence, for phage fragments whose length is shorter than Len_group (the longest length of their group), we used [0,0,0,0] to pad the end of these fragments to Len_group to make all fragments in the same group get a 4×Len_group matrix. The reverse complement sequences of all phage fragments were performed with the same operation to obtain other 4×Len_group matrices. These two matrices were concatenated to form the ‘Input1’, which is a 4×2Len_group matrix. Then we annotated the regions of coding sequence (CDS) for each contig from the training set through GenBank annotation. It is worth noting that for the phage fragments in the test set, considering that researchers generally do not have sufficient annotation information for the query phage when using our tool, we just used the gene prediction tool Prodigal (Hyatt *et al*., 2012) to annotate the CDS region of phage fragments from the test set. The CDS information of each fragment was also represented by “one-hot” encoding form, vector [1,0] and [0,1] indicates noncoding region and coding region, respectively. As before, [0,0] was used to pad this matrix and ‘Input2’ is a 2×2Len_group matrix. For prokaryotes, i.e. candidate hosts, we first clustered the prokaryote genomes by their genera, then we calculated the di-codon frequency of CDS and the 5-mer frequency of all prokaryote genomes in each cluster of different genera. The di-codon frequency of a specific genus (‘Input3’) is a 64 × 64 matrix and the 5-mer frequency (‘Input4’) is a 1024-dimensional vector. Finally, we constructed the pair of each phage fragment and each candidate host genus and assigned a label to it according to the annotations of Virus-Host DB. Label 0 indicates that the pair has no infectious relationship while label 1 means that there is an infectious annotation between the phage and a prokaryote belonging to the genus in VirusHost DB.

**Fig. 1.**
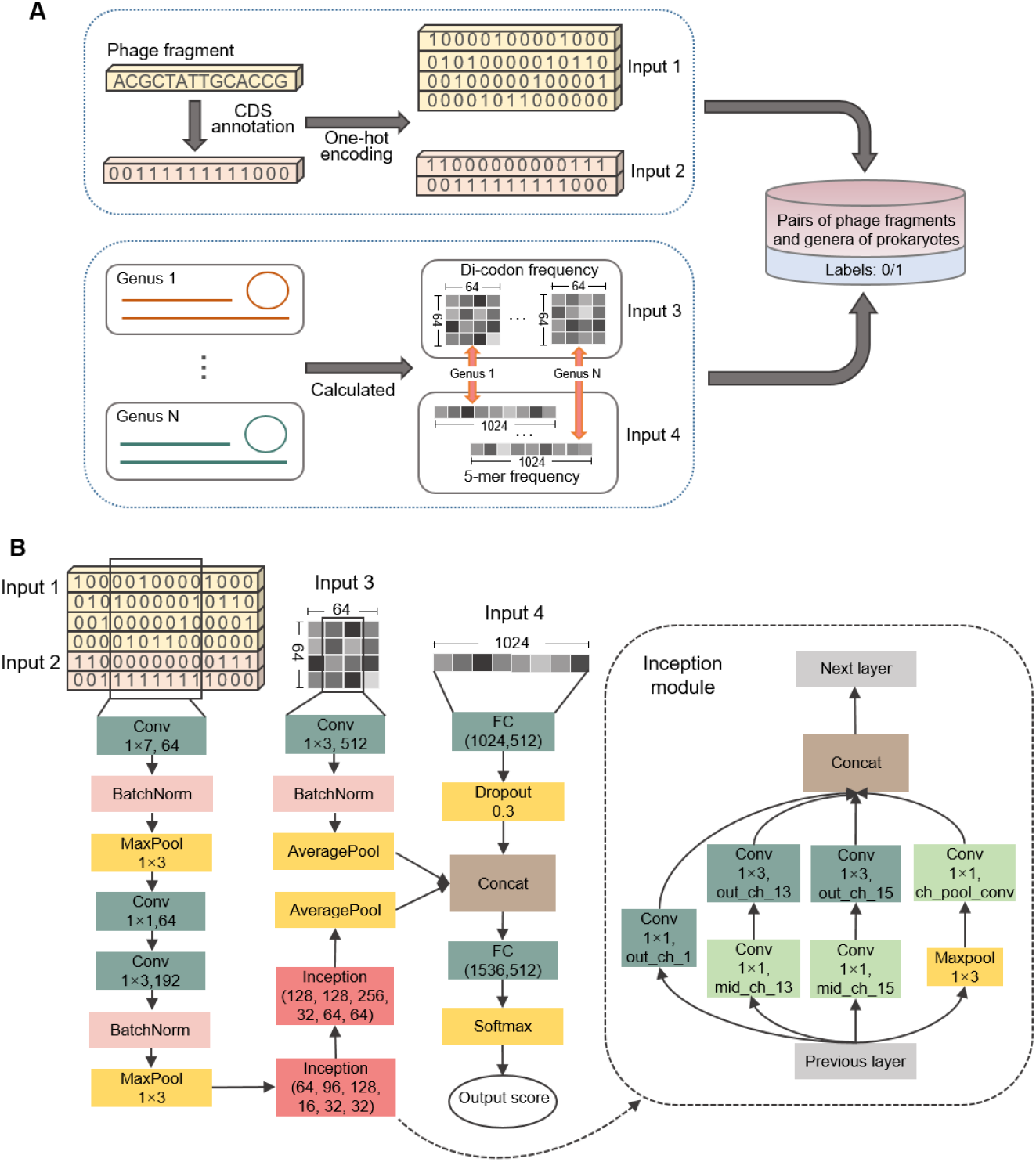
Data preparation for HoPhage-G and structure of deep learning neural networks in HoPhage-G. A). Constructing pairs of phage fragments and genera of prokaryotes and assigning labels to the pairs. B). Conv: Convolution neural network layer, BatchNorm: Batch normalization, FC: Fully connected layer. The numbers in Conv layer (green box) are kernel size and the number of channels. The numbers in FC layer (green box) are the input size and the output size. The six numbers in Inception module (red box) corresponding to out_ch_1, mid_ch_13, out_ch_13, mid_ch_15, out_ch_15, ch_pool_conv in dashed box on the right.

HoPhage-G module was built up by a deep learning model based on the inception module adopted from GoolgLeNet (Szegedy *et al*., 2016) and the normal convolution neural network. The structure of the deep learning model of HoPhage-G is shown in Fig. 1B. The inception module is also based on the convolution neural network, and it innovatively adopts a multi-path design, and each path uses a different convolution kernel size. Therefore, sequence features can be extracted at multiple scales by using the inception module. For each pair, HoPhage-G takes four inputs described above and outputs a score representing the possibility that the phage fragment and a prokaryote belonging to the genus has an infectious relationship.

Since the input length of the deep learning model is fixed, we independently trained the deep learning model in HoPhage-G for the three groups mentioned above. Due to the training set and the test set were separated before the phage fragments were simulated, we just respectively used the pairs constructed from genomes in the training set and the test set to train and test the model in HoPhage-G. However, a phage fragment often forms only one positive pair with one of the candidate host genera, and all other combinations are negative pairs. For the pairs used to train models in HoPhage-G, it is easy to think of an imbalance between the positive and negative samples. When there were a large number of candidate genera of hosts, this imbalance would be more serious. In our prokaryotes dataset, 1192 genera were included. To overcome this problem, we only constructed positive and negative pairs within one mini-batch during the training process. The size of the mini-batch was set as 8, this means that the ratio of positive pairs to negative pairs during the training process has changed from 1:1192 to 1:8. We further selected another 4 genera that do not exist in the minibatch to construct some additional negative pairs. These processes much help train the deep learning model. Moreover, to alleviate potential false-negative interactions, we only constructed negative pairs on the phage fragments and the genera which do not belong to the same family as the annotated host of this phage. But we still used all candidate 1192 genera to construct pairs for each phage fragment in the test set because it is unrealistic to correctly reduce the candidate host range to 8 genera in advance. During the training process, one-tenth of the training data was randomly selected as the validation set to tune the parameters in the deep learning model and determine when to stop training. After the parameters were adjusted by the validation set, all data in the training set was used to retrain the model in HoPhage-G with the same parameters.

However, it is cannot be ignored that phages with host annotation are relatively limited and many available phage genomes are distributed over a small proportion of the prokaryote genera in the current database. Since the machine learning algorithm relies much on the training dataset, it is difficult for any machine learning algorithm to handle totally novel data without any known knowledge related to it. So we could anticipate that for the results of HoPhage-G, compared with some dominant host genera, some of the genera which just have few phages annotated to infect them would have a poorer prediction performance.

### 2.3. HoPhage-S: the codon Markov chain model based module

To solve the above-mentioned problem that some of the genera with a small amount of data may obtain a poor prediction performance, we developed an auxiliary module HoPhage-S. Because of the genome amelioration, foreign DNA will change its sequence signatures toward its host (Suzuki *et al*., 2010). The Markov chain model has been widely used to measure the sequence signatures similarity between two DNA sequences. In previous work, WIsH used the Markov chain as a mathematical model to predict the host of phage and the ability of WIsH in predicting hosts of phages was further verified by VirHostMatcher-Net since it integrated the score of WIsH to improve performance. However, the Markov chain model in WIsH was constructed based on the base sequence. It has been pointed that most phages have to adapt to the tRNA pool of their host due to the lack of tRNA and show more consistent codon usage biases with their host (Carbone 2008). Besides, our related work showed that constructing the mathematical model based on the codon sequence was more effective than on base sequence because the sequence signatures were more significant in the coding region (Fang *et al*., 2019). Considering that the CDS density of phage is much higher than that of bacteria (Amgarten *et al*., 2018) and the reasons mentioned above, in HoPhage-S, we trained a homogeneous codon Markov chain model for each candidate prokaryote genome using codon sequences of the CDS in it and calculated the log-likelihood of a phage fragment based on each codon Markov chain model. This module was named HoPhage-S because the Markov chain model was constructed for a single prokaryote genome (strain).

For each prokaryote in our candidate host dataset, the CDSs of each prokaryote genome were extracted based on the annotation information in GeneBank. We assumed CDS^i^_n_, represents the n^th^ CDS region on prokaryote i; N^i^ represents the total number of CDSs in prokaryote i; x_1_x_2_…x_k_ represents a codon sequence; #x_1_x_2_…x_k_ represents the number of this specific codon sequence in a certain CDS region. Then the transition probability of the codon Markov chain model for prokaryote i can be represented as followed:

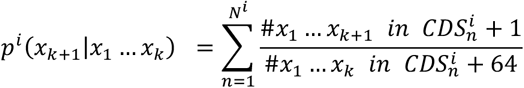

where we set k=2 in HoPhage-S, so the transition probability of the Markov chain model is a 4096 × 64 matrix.

For a given phage DNA fragment to be predicted, the CDS regions were firstly extracted using Prodigal (Hyatt *et al*., 2012). Then the similarity score of codon sequence signatures of this query fragment which based on the codon Markov chain model of prokaryote i was calculated as:

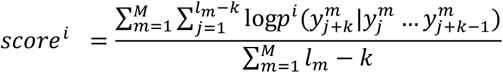

where y^m^_j_ represents the j^th^ codon of the m^th^ CDS region on the phage fragment, M represents the total number of CDS regions on the fragment, l_m_ represents the number of codons on the m^th^ CDS.

When the query phage fragment is too short, in a few cases, no CDS on the fragment could be annotated by Prodigal. This might be because the gene prediction software misses some positive predictions, or there is indeed no CDS on the given fragment. In this case, we would directly extract six codon sequences from the six phases of this phage DNA fragment, and each codon sequence would be served as a CDS to calculate the similarity score. The codon sequence with the maximum score would be considered as the *score^i^*.

### 2.4. Module integration

The whole workflow of HoPhage is shown in Fig. 2. For a query phage fragment, the CDS regions are firstly annotated by Prodigal. Then HoPhage employs HoPhage-G to score the pairs of this phage fragment and each candidate host genus in the database.

**Fig. 2.**
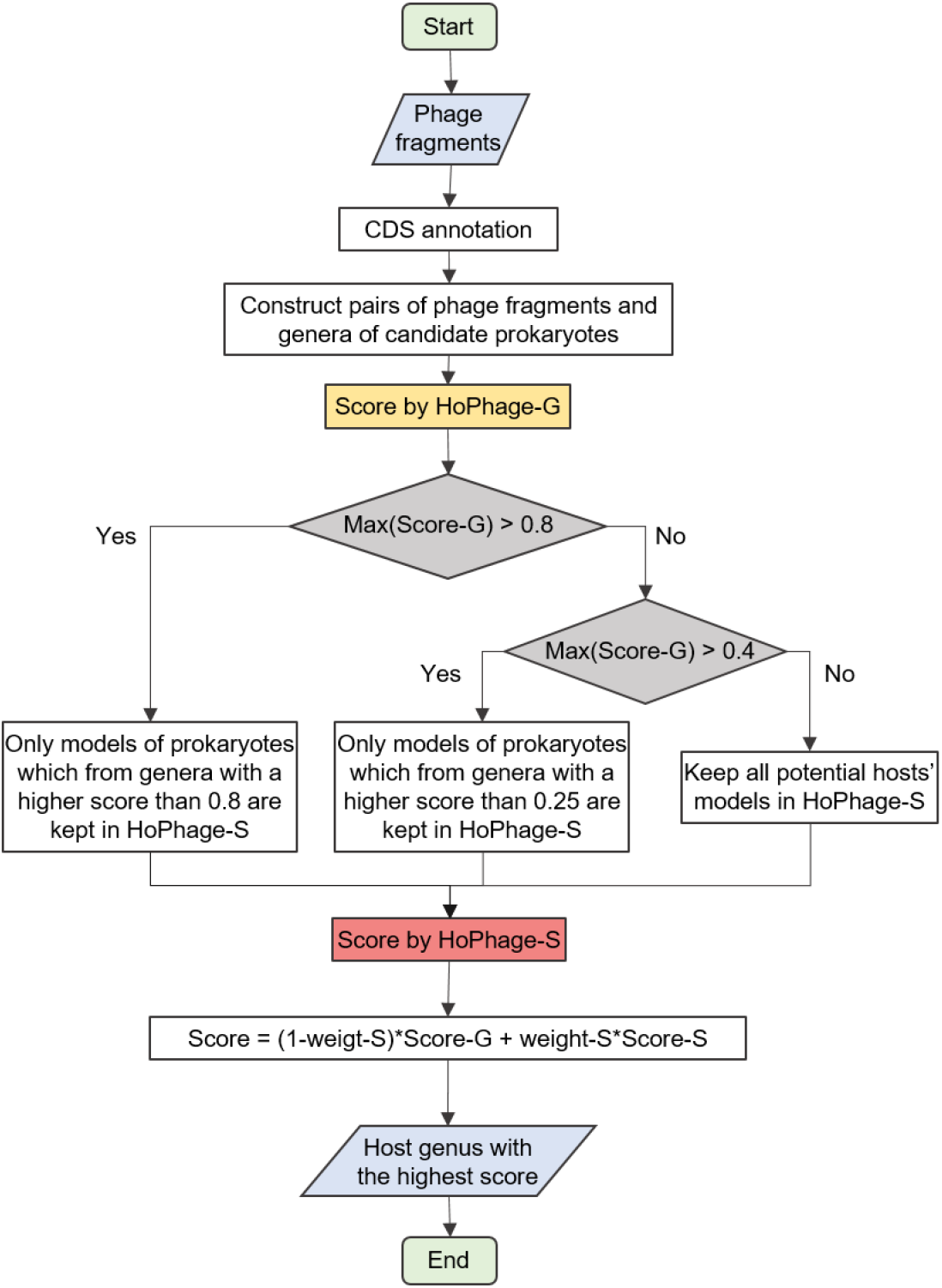
The flowchart of HoPhage. HoPhage-G is first used to predict the host of given phage fragments, then how to use HoPhage-S depends on the score of HoPhage-G.

For a binary classification task, the output of the deep learning model can represent the possibility that it is a positive sample. 0.5 is the default threshold for the prediction. A higher score means that it is more likely to be a positive sample, and it also means that the prediction is more reliable. Hence, pairs of phage fragments and candidate host genera are divided into three categories depending on the score of HoPhage-G and then different strategies are used to predict the host:

1. The highest score of all pairs in HoPhage-G (i.e. Score_Gmax) is greater than 0.8. In this case, the results of HoPhage-G are highly reliable, so in HoPhage-S we only retain the Markov chain models of prokaryotes which belong to the genera with a score higher than 0.8
2. The Score_Gmax is between 0.4~0.8 In this case, the reliability of the results of HoPhage-G is relatively high, so in HoPhage-S we retain the Markov chain models of prokaryotes which belong to the genera with a score higher than 0.25 in HoPhage-G.
3. The Score_Gmax is less than 0.4. In this case, the reliability of the score of HoPhage-G’s deep learning model is low. Therefore, in HoPhage-S we retain the Markov chain models of all prokaryotes in the existing data set.

Then the codon Markov chain models of prokaryotes retained in HoPhage-S are used to score each phage fragment, the maximum score among all Markov chain models constructed from prokaryotes that belong to the same genus will represent the score of this genus. All scores obtained by HoPhage-S are normalized to [0, Score_Gmax]. Hence, the weighted average of the highest score among one prokaryote genus in HoPhage-S and the score of this genus in HoPhage-G is set as the incorporated score for this genus. Finally, the genus with the highest score is used as the default output of HoPhage, indicating that this phage fragment is most likely to infect prokaryotes belonging to this genus.

## 3. Results

### 3.1 Evaluation of the performance on artificial phage fragments

To evaluate the prediction performance of HoPhage on short phage fragments, the benchmark datasets of artificial short contigs with three different lengths, 100-400 bp, 401-800 bp and 801-1,200 bp, were generated.

We first assessed the performance of host prediction using HoPhage-G and HoPhage-S individuals alone. Results showed that HoPhage-G outperforms other existing tools and achieved AUCs (area under ROC curve) of 0.989~0.993, which were evidently higher than that of other tools (Fig. S1B). As for the HoPhage-S, the AUCs of it were 0.712~0.739, which were also better than these tools in general (Fig. S3). More detailed results and comparisons with other tools are included in the Supplementary Material.

Considering that in practical applications, there are not as many as 1192 prokaryotic genera dominant in a microbial community, we limited the range of candidate hosts to 50 genera. For each comparison, we randomly selected 50 host genera from 155 genera included in Virus-Host DB as candidate host genera. Once the range of host genera is clarified, we would only use these genera and phage fragments to form the pairs in HoPhage-G, and only keep models constructed by the prokaryotic genomes belonging to these genera in HoPhage-S, and test fragments generated from phage genomes that can infect prokaryotes from one of these 50 genera were retained to evaluate model performance. Such an evaluation was repeated 20 times for each group in the test sets.

We set the weight of HoPhage-G/HoPhage-S to 0.5/0.5 and compared the performance of incorporated prediction of HoPhage with other tools. The details on the weight selection are described in Supplementary Material. Although the advantages of HoPhage-S compared with other tools were not significant, after narrowing the host range by HoPhage-G in advance, HoPhage which integrates these two modules achieved significantly better performance compared with all other tools. The prediction accuracy was calculated as the percentage of phage fragments whose predicted hosts had the same taxonomy as their respective annotated hosts. As a result, the average accuracies of HoPhage for the three groups were 25.0%, 29.0%, and 31.4% higher than that of WIsH at the genus level (Fig. 3), respectively.

**Fig. 3.**
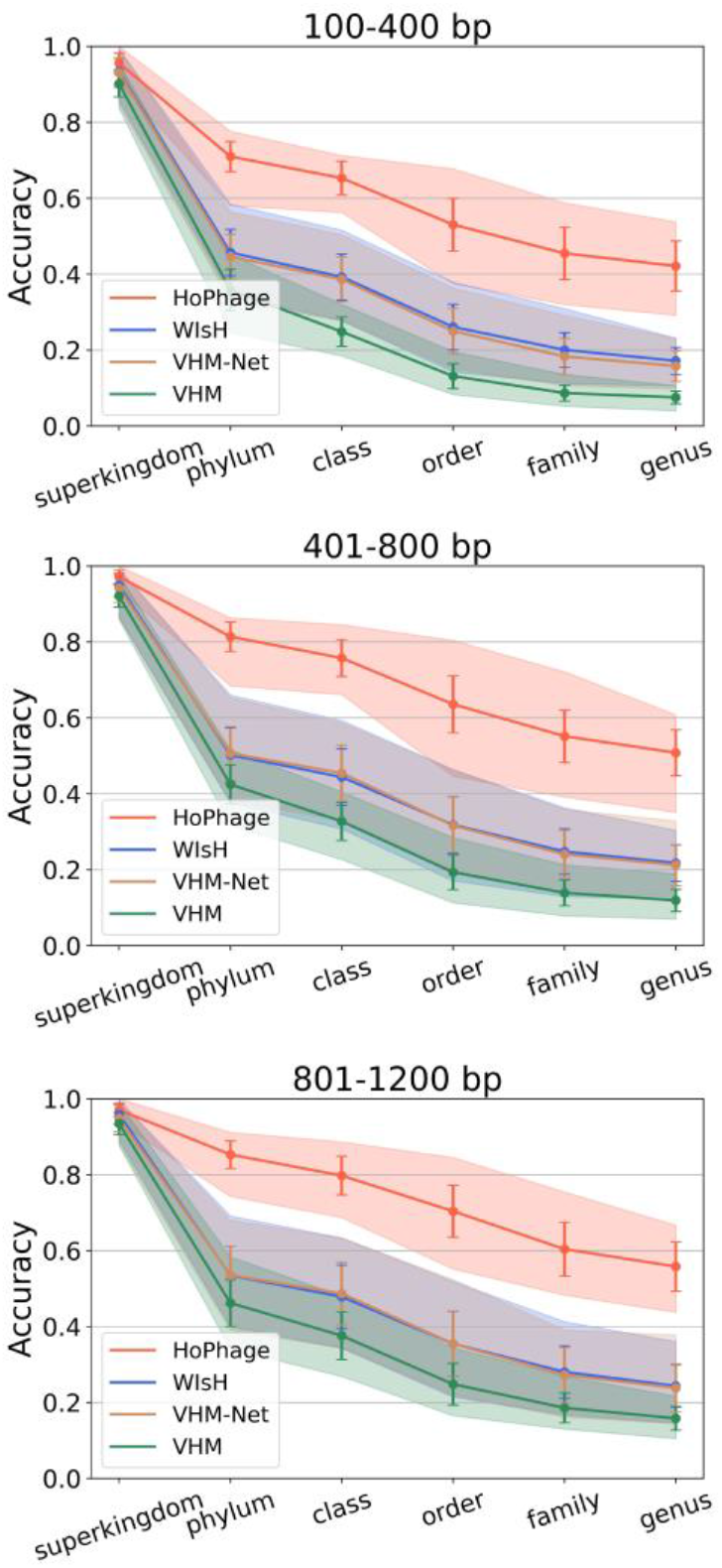
Performance of HoPhage on artificial phage fragments. Prediction accuracies of HoPhage at different taxonomic levels and comparisons with related tools. VHM-Net: VirHostMatch-er-Net, VHM: VirHostMatcher. The solid lines with error bars are the average accuracy of 20 randomly selected data. The light-colored area indicates the range of prediction accuracies.

### 3.2. Evaluation of the performance on longer fragments

We further constructed two additional length intervals, which are 1,201-3,000bp and 3,001-5,000bp, to evaluate the performance of HoPhage on longer phage fragments. As we only trained the deep learning-based model in HoPhage-G on the above three preliminary groups with the shorter input size, we first evaluated the performance of using HoPhage-G alone to test the model’s ability to handle sequences whose length is not within the preset range. For sequences longer than 1,200 bp, a scan window will move across the sequence without overlapping, and the weighted average score of all windows’ predictions is calculated. Confusion matrics (Fig. 4) of these two groups showed that HoPhage-G achieved better performance on longer phage fragments, despite using the models training by shorter fragments. The final prediction accuracies (Fig. 5) of HoPhage at every taxonomic level were also much better than related tools.

**Fig. 4.**
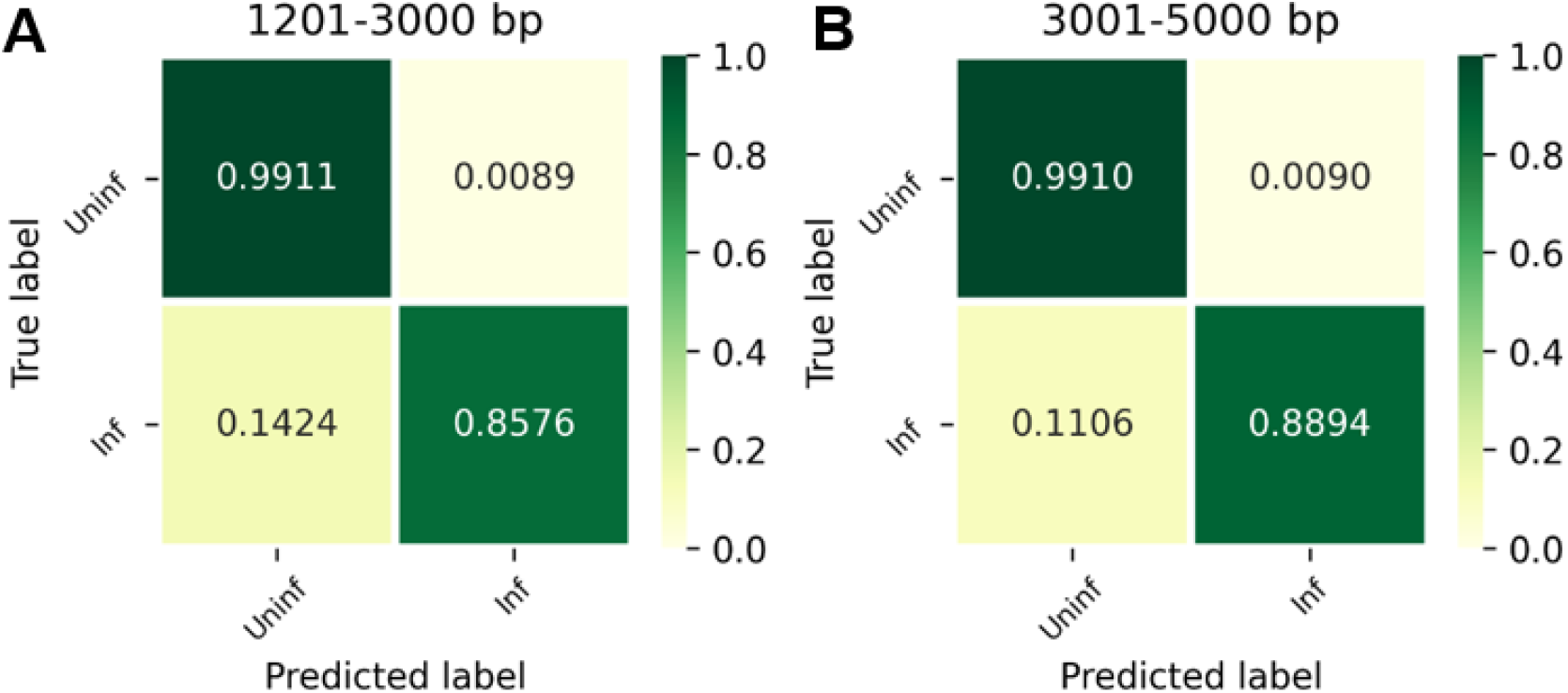
Confusion matrices of HoPhage-G on longer phage fragments. ‘Inf’ means that the pair of phage fragment and host genus has an infection relationship while ‘Uninf’ means that there is no infection relationship.

**Fig. 5.**
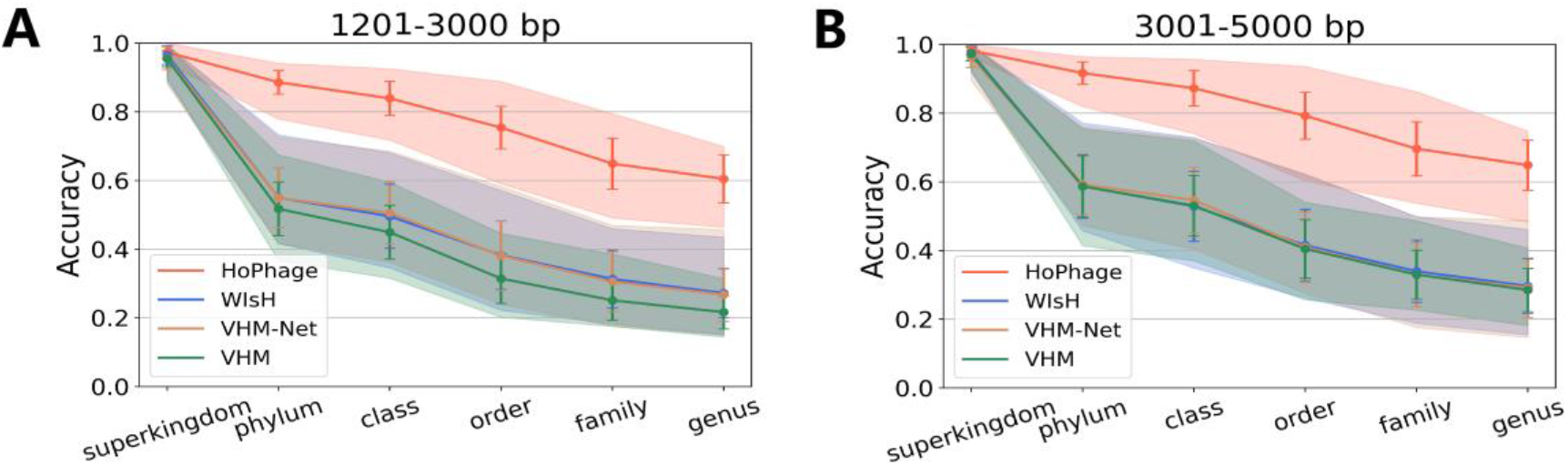
Prediction accuracies of HoPhage on longer phage fragments and comparison with the related tools. A, B are the accuracies of the top 1 prediction of host genus of HoPhage-S, WIsH, VHM-Net and VHM on 1200-3000, 3,000-5,000, respectively. Orange, blue, brown, green lines represent the results of HoPhage, WIsH, VHM-Net and VHM, respectively. The solid lines with error bars are the average accuracy of 20 randomly selected data. The light-colored area indicates the range of prediction accuracies.

### 3.3. Evaluation of the performance on real data from the mock virus communities

We also used the real virome data set to evaluate the host prediction performance of HoPhage. This data set comes from the mock virus communities which are comprised of 12 specific phages that grow on *Pseudoalteromonas, Cellulophaga baltica,* and *Escherichia coli* (Roux *et al*., 2016). The virus particles were enriched together and sequenced. We downloaded the assembled contigs from three samples, MCB1, MCB2, and MCB3. Then we generated a local *BLAST* database with 12 phage genomes mentioned above and used *blastn* to trace back the source of the phage contigs, fragments with e-value less than 0.1 and length longer than 800 bp were kept. For the fairness of comparison, we excluded the 12 phages contained in the mock virus communities and regenerated the training data, and then retrained the deep learning model in module HoPhage-G.

Among the host range of all 1192 genera in our data set, the overall accuracy of HoPhage at the genus level was 70.90% while WIsH was 58.54%, VirHostMatcher-Net was 42.70%. For each sample, the host prediction accuracies of HoPhage, WIsH and VirHostMatcher-Net of these contigs at the genus level are shown in Fig. 6. The average host prediction accuracies of HoPhage for phage contigs of three samples whose hosts are *Cellulophaga, Pseudoalteromonas,* and *Escherichia* were 34.85%, 84.44%, 44.44%, respectively, which were 1.89%, 12.54%, and 44.44% higher than those of WIsH and 32.20%, 24.62%, 44.44% higher than those of VirHostMatcher-Net. In addition, WIsH and VirHostMatcher-Net did not correctly predict any phage contig whose host belonging to *Escherichia*. These results on the real virome data showed that HoPhage does have obvious advantages in predicting hosts of phage contigs.

**Fig. 6.**
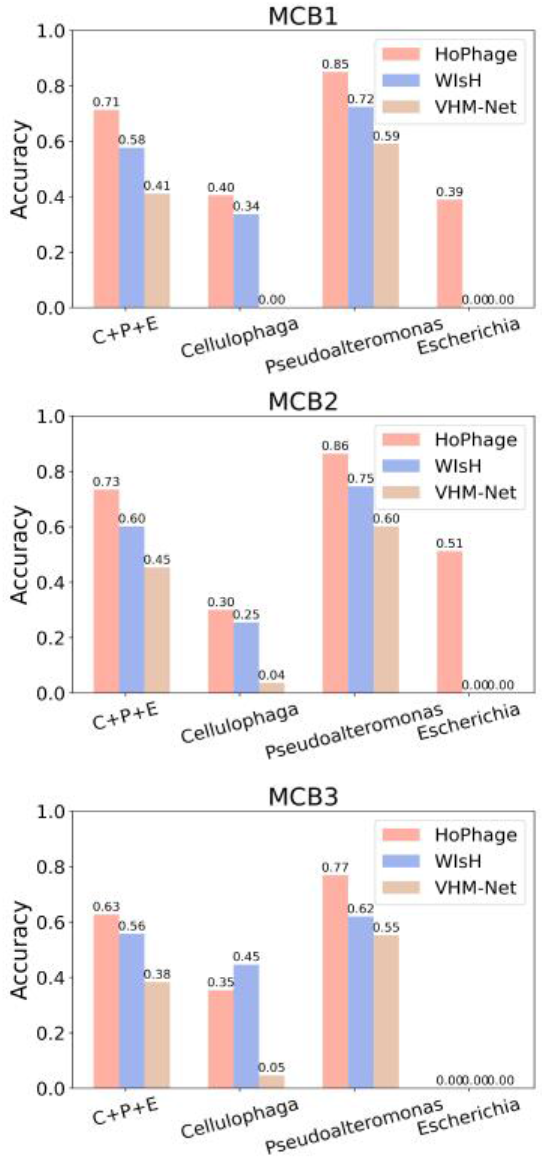
Genus accuracies of HoPhage and related tools on contigs from three real virome samples. ‘C+P+E’ indicates the overall accuracy of all three genera, while ‘Cellulophaga’, ‘Pseudoalteromonas’, and ‘Escherichia’ are calculated separately.

In this practical application, we adjusted the scoring weights of the two modules in HoPhage according to the preliminary results. The above statistical results were finally obtained by setting the weight of HoPhage-G as 0.2 and the weight of HoPhage-S as 0.8. In this process, we found that increasing the weight of HoPhage-G can improve the prediction performance of phage contigs whose host is *Escherichia*, but at the same time, the prediction accuracy of phage contigs whose host is the other two genera decreases. When increasing the weight of HoPhage-S, the situation was just the opposite. This is probably because the volume of phages that can infect *Escherichia* is large, so that the deep learning model in HoPhage-G can well summarize the relationships between these phages fragments and potential hosts and make better predictions, while the other two genera cannot obtain good prediction in deep learning model due to the relative lack of related data. Therefore, if possible, we recommend users to choose appropriate weights for the two modules in HoPhage based on the community from which the phage fragments come when using HoPhage. If there are many dominant genera in this community belonging to the categories which have a large number of related records, the higher weight of module HoPhage-G may improve the prediction performance as a whole.

### 3.4. Exploration of the marker genes in phages through HoPhage-S

It has been pointed out that the evolutionary pressure of phage genome in the co-evolution process with hosts is not uniform, and the codon usage preference will only be more prominent on some genes of phages (Carbone 2008). Since phages lack conservative genes like 16S rRNA in prokaryotes, the taxonomic classification of phages often depends on their morphology. Therefore, we assumed that based on the potential of a phage gene in identifying its host, genes that are more consistent with their host during the co-evolution can be regarded as the marker genes of phages.

As can be seen from our above results, HoPhage-S, which used the codon Markov chain model to measure the similarity between phage fragments and potential hosts, showed better performance than WIsH which used the base Markov chain model. In order to quantify the potential of different genes of phage in identifying the host, we used all single genes extracted from the phages in the training and test set as the inputs of HoPhage-S for host prediction. Then we calculated the accuracies of host prediction at the genus level for genes with the annotations containing different keywords. 16 keywords were used to extract the DNA sequence of annotated genes from phage genomes. According to the functions of the proteins they encode, these genes were divided into three categories, including proteins used for the processes of the central dogma (‘polymerase’, ‘ligase’, ‘primase’, ‘helicase’, ‘exonuclease’), proteins function in morphology (‘head’, ‘tail’, ‘scaffold’, ‘capsid’, ‘fiber’, ‘baseplate’) and proteins involved in the infection processes (‘lysin’, ‘holin’, ‘integrase’, ‘transposase’, ‘excisionase’).

We used the ratio of prediction accuracy at the genus level to gene length to quantify the potential of a gene to identify the host and used the potential of genes whose annotations include “polymerase” as the reference value 1. As shown in Table 1 and Fig. 7, there was roughly a trend that infection-related genes had the greatest potential, morphology-related genes were the second, and the central dogma-related genes had the lowest potential. In more detail, in addition to the two genes encoding non-important structural proteins, baseplate and fiber, other morphology-related genes were slightly more potential to identify hosts than the central dogma-related genes, and the potential of infection-related genes was significantly higher than the previous two categories. This result was reasonable and in line with our expectations since the proteins involved in infection processes can directly interact with the host, which will inevitably lead to more evolutionary pressure of these relevant genes from their hosts in the co-evolution. It is worth noting that the functions of proteins ‘integrase’, ‘transposase’ and ‘excisionase’ are integration genome sequence of temperate phage into or excision it from the host chromosome (Baker 1995; Cho *et al*., 2002), while proteins ‘holin’ and ‘lysis’ are function in cytolytic lysis process (Ugorcakova *et al*., 2003) which exists in both temperate and virulent phages. The results showed that genes encoding temperate phage-specific proteins had a higher potential for host identification. For example, the ‘excisionase’ gene achieved a 51.61% accuracy of host prediction at the genus level as the average length of this gene was as short as 287 bp, hence its potential in identifying the host was 18 times that of gene ‘polymerase’. This was also rational since temperate phages may integrate their genome into the host chromosome and reproduce with the host, therefore, adapting more host sequence signatures for better survival.

**Table 1.** Prediction accuracy of HoPhage at the genus level with different weights.

| keyword | accuracy | average length of genes (bp) | number of genes | potential |
| --- | --- | --- | --- | --- |
| **polymerase** | 16.54% | 1683 | 3423 | 1 |
| **ligase** | 11.17% | 1106 | 1101 | 1.027653 |
| **primase** | 13.59% | 1231 | 1854 | 1.123337 |
| **helicase** | 17.57% | 1402 | 2971 | 1.275183 |
| **exonuclease** | 15.94% | 903 | 1286 | 1.796177 |
| **baseplate** | 8.09% | 1225 | 2956 | 0.671987 |
| **fiber** | 14.20% | 1785 | 2521 | 0.809466 |
| **tail** | 21.69% | 1242 | 14427 | 1.776996 |
| **capsid** | 25.84% | 1096 | 3285 | 2.399002 |
| **head** | 17.52% | 711 | 4367 | 2.507339 |
| **scaffold** | 17.80% | 712 | 1337 | 2.543833 |
| **lysin** | 30.46% | 1099 | 2088 | 2.820206 |
| **holin** | **27.69%** | **364** | **1477** | **7.740521** |
| **integrase** | 50.72% | 1153 | 905 | 4.476087 |
| **transposase** | 51.62% | 912 | 277 | 5.759327 |
| **excisionase** | **51.61%** | **287** | **93** | **18.29787** |

**Fig. 7.**
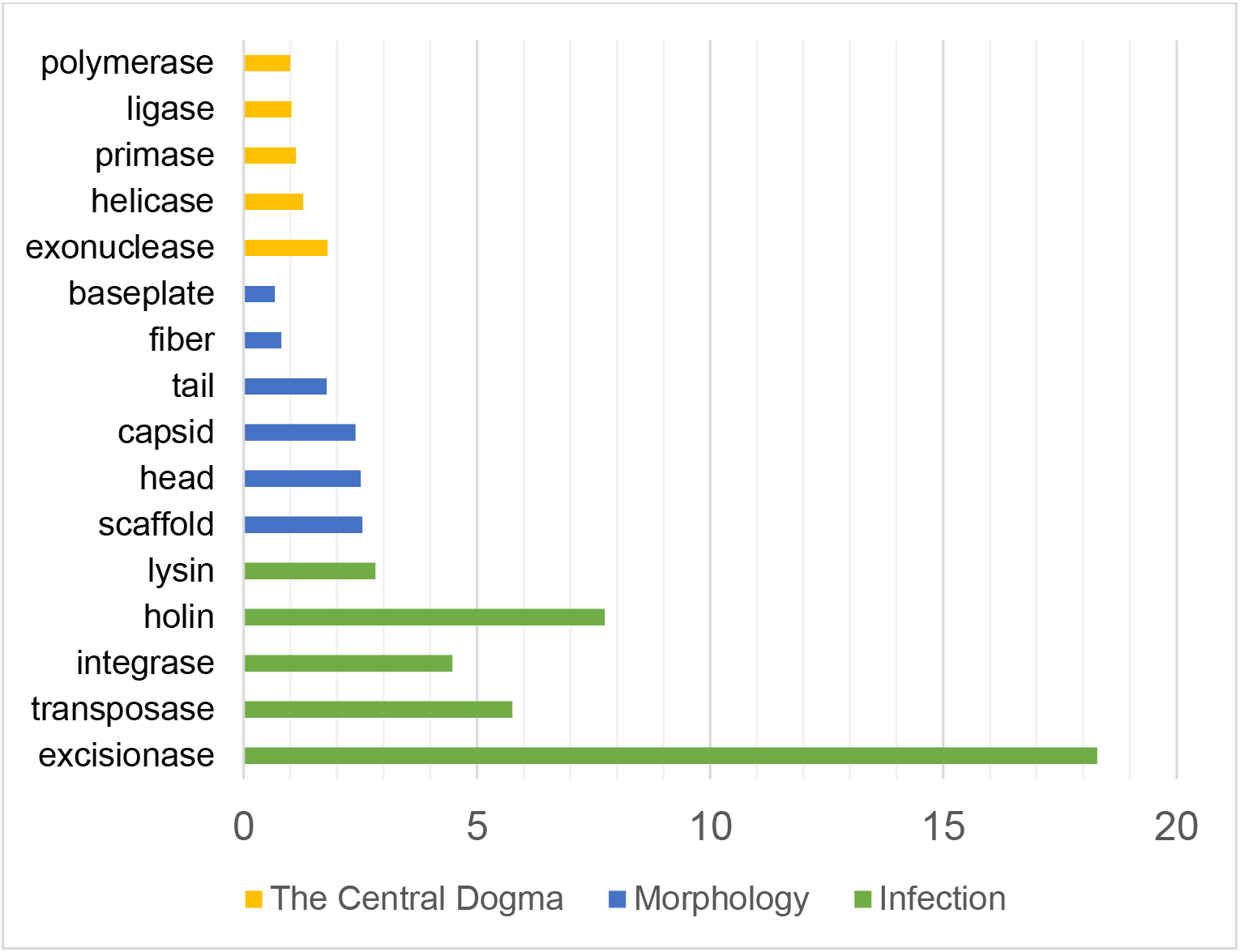
The potential of phage genes annotated with different keywords in identifying host.

The tetranucleotide frequency has been used to construct a phylogenetic tree of phages and found that phages with the same host converge on the tree (Pride *et al*., 2006). As holin exists in both temperate and virulent phages, we further constructed the phylogenetic tree of phages using the genes annotated as ‘holin’ or ‘Holin’. The holin genes which correctly predicted the host genus and gained scores ranked in the top 50 were selected. Fig. 8 is the phylogenetic tree constructed by these 50 holin genes. It can be seen that genes derived from phages with the same host converge on the tree. We believed that the genes which have high potential in host identification can be used as phage marker genes for an alternative taxonomic classification of phages.

**Fig. 8.**
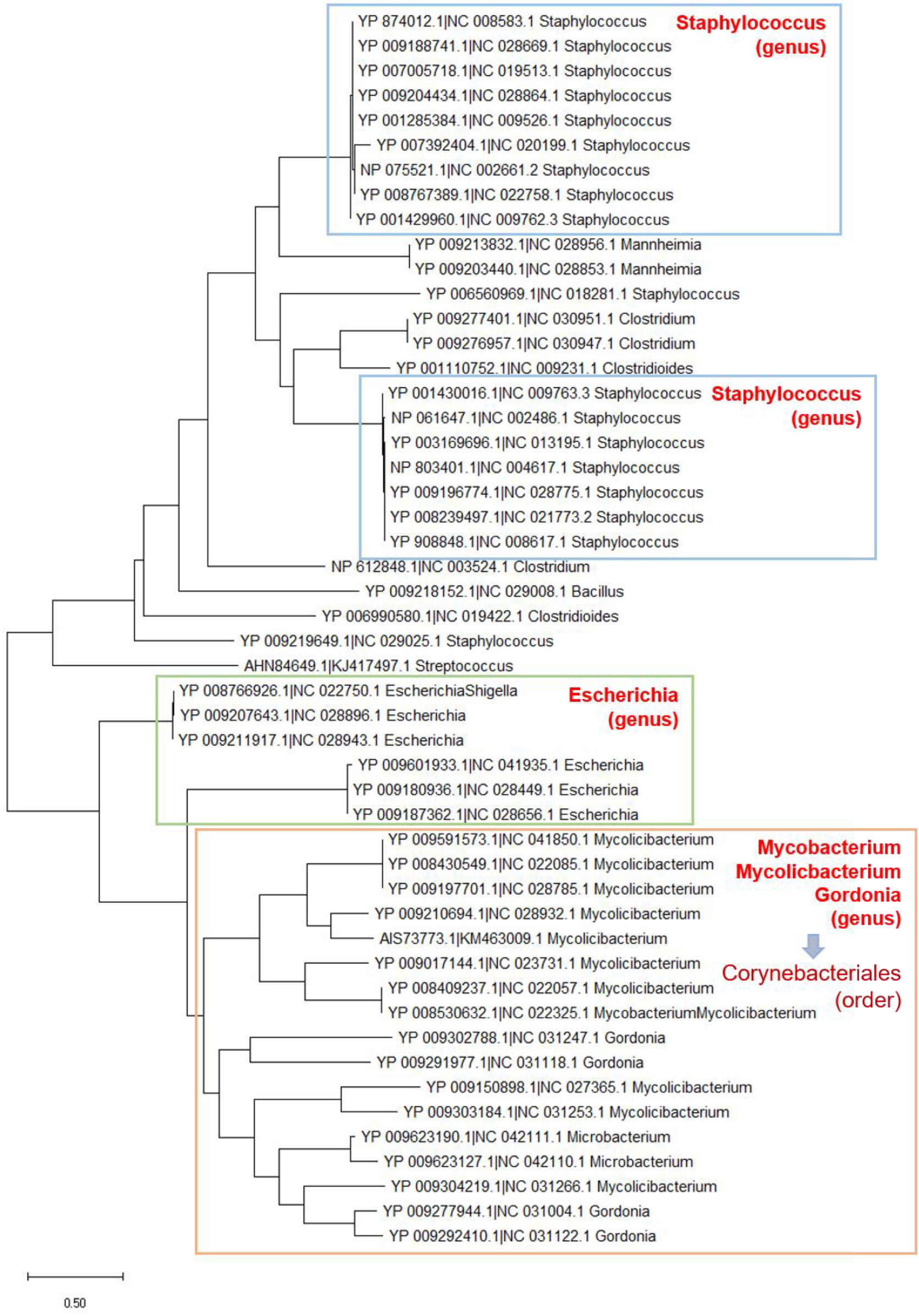
Phylogenetic tree constructed by the DNA sequence of ‘holin’ genes from phages.

## Discussion

In this paper, we present HoPhage, an ab initio tool for identifying hosts of phage fragments from metavirome using the sequence signatures. Testing on a benchmark dataset of artificial short contigs and real virome data of mock viral community shows that HoPhage performs much better than the state-of-the-art tools for short fragments within a wide candidate host range. HoPhage can directly be employed on virome data, in which the viral particle is enriched before sequencing. For untargeted metagenomic data, users need to firstly identify the phage contigs from chromosome-derived contigs using related software such as VirSorter (Roux *et al*., 2015), VirFinder (Ren *et al*., 2017), PPR-Meta (Fang *et al*., 2019), and DeepVirFinder (Ren *et al*., 2020) and then use HoPhage to identify the host of the phage fragments. Like other host prediction software, users can specify the candidate host for HoPhage. In the released package, HoPhage-G contains pre-calculated di-codon frequencies and 5-mer frequencies of 1192 genera from 5314 prokaryotes and HoPhage-S contains all Markov chain models trained by these 5314 prokaryotes. Under default parameters, HoPhage will use all candidate hosts in the dataset. In addition, users can also provide a list to restrict the host range for virome data from a certain environment. For example, in the human gut, about 15 bacterial genera occupy a total of 70% of organisms of the microbial community (Li *et al*., 2014). When employing HoPhage over gut virome data, users can restrict the candidate host within these genera to avoid false-positive prediction of a host that does not exist in this environment. On the other hand, our evaluations show that HoPhage can also achieve satisfactory performance even if there are hundreds of candidate host genera, which means that the lack of prior knowledge about the candidate host range will not serious affect the usage of HoPhage.

To make a reliable prediction, we designed two modules to improve HoPhage’s performance, named HoPhage-G and HoPhage-S. HoPhage-G is a deep learning-based module. Through constructing pairs of phage fragments and genera of potential hosts, this complex multi-classification host prediction issue transforms into a two-classification task. Hence HoPhage-G aims to judge whether the query phage fragment can infect a prokaryote from a specific genus. We also adopt the inception module from GoogLeNet, which can extract features at multiple scales. While deep learning algorithms have shown a strong ability to extract sequence signatures over a large scale dataset to make a reliable prediction and have already been employed by many tools to predict the interaction between biological components, HoPhage-G demonstrates superior performance as it can obtain a high TPR under a very low FPR. However, machine learning-based tools rely on existing data that is used for training. Since the distribution of the number of phages that can infect hosts from different genera in the current database is unbalanced and a large number of known phages derives from a narrow host range, therefore HoPhage-G presents a poorer performance for some phages which lack the related data. So we designed HoPhage-S as the complement, which is a Markov chain-based module to overcome this unbalanced problem. The innovation of HoPhage-S is that we employed a codon Markov chain model for CDS regions in the prokaryote genomes, rather than the base Markov chain model that WIsH employed. It has been shown that sequence signatures are more concentrated in the coding sequence and the density of CDS on the phage genome is higher than that in the prokaryote genome. Our results show that the codon Markov chain model is indeed more effective than the base Markov chain model. At last, testing on the artificial benchmark dataset of artificial phage contigs and real virome data, HoPhage demonstrates much better performance on short fragments within a wide candidate host range at every taxonomic level.

However, there are still some shortcomings in our work and needed to solve in the future. It must be admitted that although HoPhage can handle novel phages, due to the limitations of the algorithm and currently accessible dataset, phages with very little relevant data will inevitably get poorer prediction accuracy. In addition to sequence signatures, it is also worth considering employing other signals to further improve the performance of HoPhage in future researches, such as the presence of CRISPR spacers or the abundance profiles. Moreover, HoPhage is designed primarily for the prokaryotic virus (i.e. phages), which is dominant in the microbial community, but the real virome data may contain a small number of eukaryotic viruses. Recently, the host prediction tool for several specific eukaryotic viruses has been designed (Mock *et al*., 2020). In order to let HoPhage more versatile, it is also worth considering the host prediction for eukaryotic virus fragments. But while most of the eukaryotic viruses are RNA viruses, which will not appear in large amounts in DNA sequencing data, this problem has little impact on the application of HoPhage. Another problem is that even in the virome data, there will still be some host contamination, hence the de-hosting operation before using HoPhage will be more conducive.

In conclusion, HoPhage demonstrates much better performance on short fragments within a much wider candidate host range. We expect HoPhage to play a vital role in identifying hosts of novel phages and help researchers to explore the underlying ecological impact of phage in a community.

## Availability of supporting data and materials

The artificial contigs, related scripts, and original results are available at http://cqb.pku.edu.cn/ZhuLab/HoPhage/data/. All the other data are available at corresponding references mentioned in the main text.

HoPhage is user-friendly and does not have high hardware requirements. We have released the program as a Docker image (https://hub.docker.com/repository/docker/jietan95/hophage) so that non-computer professionals can use HoPhage without installing any dependency package. Besides, the physical host version of HoPhage can speed up with GPU and is more suitable to handle large-scale data. The program is freely available at http://cqb.pku.edu.cn/ZhuLab/HoPhage/ or https://github.com/jie-tan/HoPhage/.

