## Supplementary Material for "Identify phage hosts from metaviromic short reads based on deep learning and Markov chain model"

### 1. The performance of using HoPhage-G alone

We first evaluated the performance of HoPhage-G alone with a very wide host range and all 1192 genera were used as the candidate host genera. The confusion matrices of three preliminary groups with different length intervals are shown in Fig. S1A. The upper right corner of the confusion matrix is the false positive rate (FPR), and the lower right corner is the true positive rate (TPR). It can be seen that HoP-G can achieve a high TPR under a very low FPR, except for the "100-400 bp" group with a relatively low TPR. To compare the performance of HoPhage-G with related tools, we regarded WIsH, VirHostMatcher-Net (VHM-Net) and VirHostMatcher (VHM) as binary classification tools too and draw ROC (Receiver Operating Characteristic) curve and PRC (Precision-Recall Curve) through the scoring of the model. The ROC curve and PRC with the value of AUC (Area Under Curve) and AP (Average Precision) are shown in Fig. S1B and Fig. S1C. HoPhage-G showed a clear superiority on all these evaluations. ROC curves are commonly used to present results for binary decision problems in machine learning. The researchers have found that for the same classifier whose test set had a balanced 1:1 class distribution, when the number of negative instances has been increased 10-fold, the ROC curve almost maintained its original performance while the PRC significantly decreased (Fawcett, 2006). For HoPhage-G, the ratio of positive and negative samples is less than 1:1000. So it can be envisioned that HoPhage-G will get a poor PRC. It also indicated that when dealing with highly skewed datasets, PRC gives a more informative picture of an algorithm's performance (Davis, *et al.*, 2006). Whether using PRC or ROC, HoPhage-G performed better than other tools, further verified the effectiveness of the model.

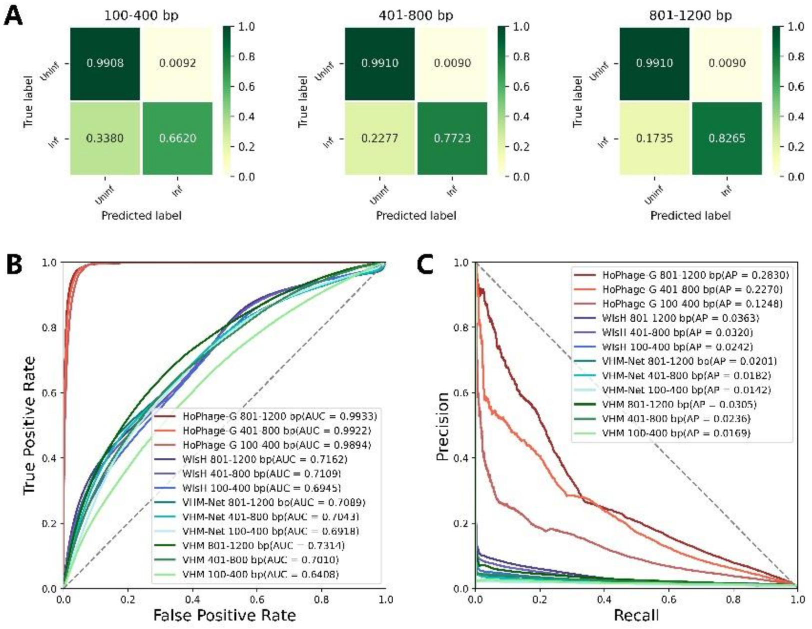


**Fig. S1. Performance of HoPhage-G and comparison with related tools**. (A) Confusion matrices of HoPhage-G. 'Inf' means that the pair of phage fragment and host genus has an infection relationship while 'Uninf' means that there is no infection relationship. (B) and (C) are the receiver operating characteristic (ROC) curve and precision-recall curve (PRC) of HoPhage-G. WIsH, VirHostMatcher-Net (VHM-Net) and VirHostMatcher (VHM) on phage fragments of different lengths, respectively. The values of area under curve (AUC) and average precision (AP) are showed in legends.

It should be pointed out that since the customized candidate hosts must exist in its existing data set when using VirHostMatcher-Net, we could not arbitrarily specify candidate hosts for it, so its host range was a bit narrower than HoPhage. In detail, 1149 genera were available in VirHostMatcher-Net while HoPhage, WIsH, VirHostMatcher using the consistent candidate hosts which range from 1192 genera. This was also the same situation in the subsequent comparisons.

Considering that in practical applications, there are not as many as 1192 prokaryotic genera dominant in a microbial community, we limited the range of candidate hosts to 50 genera. For each comparison, we randomly selected 50 host genera from 155 genera included in Virus-Host DB as candidate host genera. Once we clarified the range of host genera, in HoPhage-G we would only use these genera and phage fragments to form the pairs, in HoPhage-S we would only keep models constructed by the prokaryotic genomes belonging to these genera, and test fragments generated from phage genomes that can infect prokaryotes from one of these 50 genera are retained to evaluate model performance. Such an evaluation was repeated 20 times for each group of test sets. For VirHostMatcher-Net, several random selections would include a few genera that are not in the 50 genera. For the sake of fairness, we narrowed the host range of VirHostMatcher-Net to all other available genera, and only retained phage fragments that can infect the remaining genera to calculate the prediction accuracy. The prediction accuracy was calculated as the percentage of phage fragments whose predicted hosts had the same taxonomy as their respective annotated hosts. Fig. S2 shows the prediction accuracies of HoPhage-G and other related tools at different taxonomic levels among the host range of 50 genera. Besides HoPhage, WIsH was relatively the most effective tool, the performance of VirHostMatcher-Net was very close to WIsH, and the prediction accuracies of VirHostMatcher were much lower than that of other tools. When only the genus with the highest score was treated as the prediction result, the accuracies at the genus level of HoPhage-G were 20-30% higher than that of WIsH (Fig. S2A, S2C, S2E). In other taxon levels, HoPhage-G also achieved much higher accuracies than WIsH. When all the genera among the top 3 scores were considered, the result of HoPhage-G was improved much more than these related tools (Fig. S2B, S2D, S2F), which indicates that HoPhage-G was more capable to give the correct result a relatively higher score than related tools.

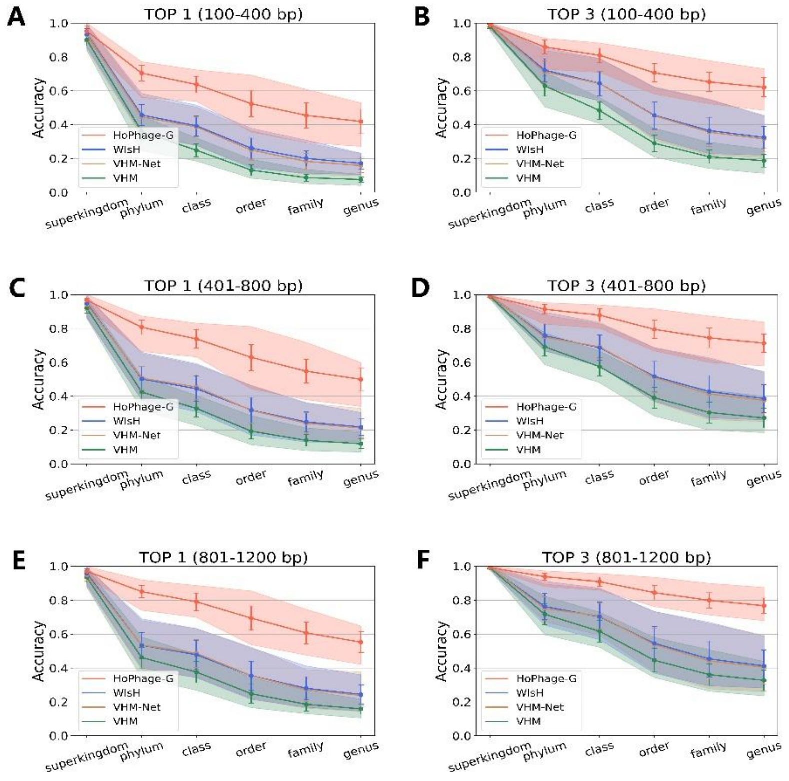


**Fig. S2. Prediction accuracy of HoPhage-G at different taxonomic levels within the host range of 50 genera and comparison with related tool.** A, C, E are the accuracies of the top 1 prediction of host genus of HoPhage-G, WIsH, VHM-Net and VHM at each taxonomic level on 100-400, 401-800, 801-1200bp, respectively. B, D, F are the top 3 predictions.

We further explored why the performance of WIsH and VirHostMatcher-Net were very close. As we mentioned earlier, when processing short phage fragments, VirHostMatcher-Net will integrate WIsH's score to predict the host. With the help of the percentile information in the outputs of VirHostMatcher-Net, users can better understand how relevant each feature score is for a particular prediction. We counted the percentile information outputted by VirHostMatcher-Net in each test set. The results indicated that the prediction of VirHostMatcher-Net was highly dependent on the score of WIsH for short phage fragments. As shown in Table S1, the distribution of the percentiles of WIsH's score and the other three features used in VirHostMatcher-Net is very extreme. Although the percentile of WIsH decreased slightly with the increase of the phage fragment length, we could still conclude that the score of WIsH plays a decisive role in the prediction of VirHostMatcher-Net for short phage fragments.

**Table S1. The distribution of the percentile values in VirHostMatcher-Net**.

| Percentile | **WIsH_pct > 0.9** | **POS_pct > 0.1** | **NEG_pct > 0.1** | **CRISPR_pct > 0.1** |
|---|---|---|---|---|
| **100-400 bp** | 99.52% | 0.12% | 0.14% | 1.58% |
| **401-800 bp** | 98.82% | 0.06% | 0.08% | 3.00% |
| **801-1200 bp** | 98.50% | 0.02% | 0.06% | 3.98% |

### 2. The performance of using HoPhage-S alone

Then we also tested the performance of using module HoPhage-S alone with the same test sets and compared it with related tools. Firstly, we also regarded HoPhage-S as a two-classifier to plot the ROC curve and calculated AUC by constructing pairs at the genus level. Compared to other existing tools, HoPhage-S also had a slight advantage (Fig. S3). The prediction accuracies of HoPhage-S at the different taxonomic levels among the host range of 50 genera were also better than these tools in general (Fig. S4). Since both HoPhage-S and WIsH are based on the Markov chain model, the difference between these two is that the former uses the codon sequence of CDS to build the Markov chain model, while the latter uses the base sequence to build the model. By comparing the performance of them, it can be concluded that whether the signatures of the codon sequence have more potential to predict the relationship between the phage and the candidate host. The details of the prediction accuracy of HoPhage-S and related tools at the genus levels among 20 randomly selected test sets are shown in Fig. S5. Although only using the information of the CDS region makes the sequence length that HoPhage-S could utilize becomes shorter, HoPhage-S still showed better performance. This result demonstrated that, for phage, the sequence signatures extracted from the codon sequence of the coding regions are indeed more similar to the sequence signatures of their host due to greater selection pressure, hence the CDS of phage has more potential to identify hosts.

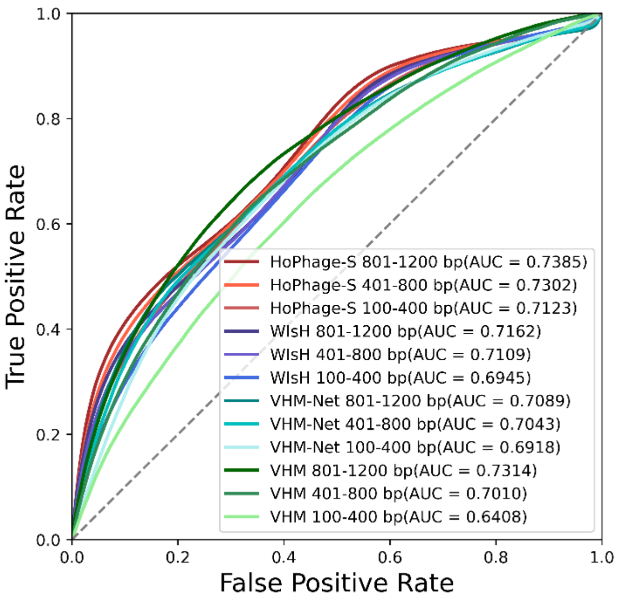


**Fig. S3. ROC of HoPhage-S and comparison with related tools**. The receiver operating characteristic (ROC) curve of HoPhage-S, WIsH, VirHostMatcher-Net (VHM-Net) and VirHostMatcher (VHM) on phage fragments of different lengths, respectively. The values of area under curve (AUC) are shown in legends.

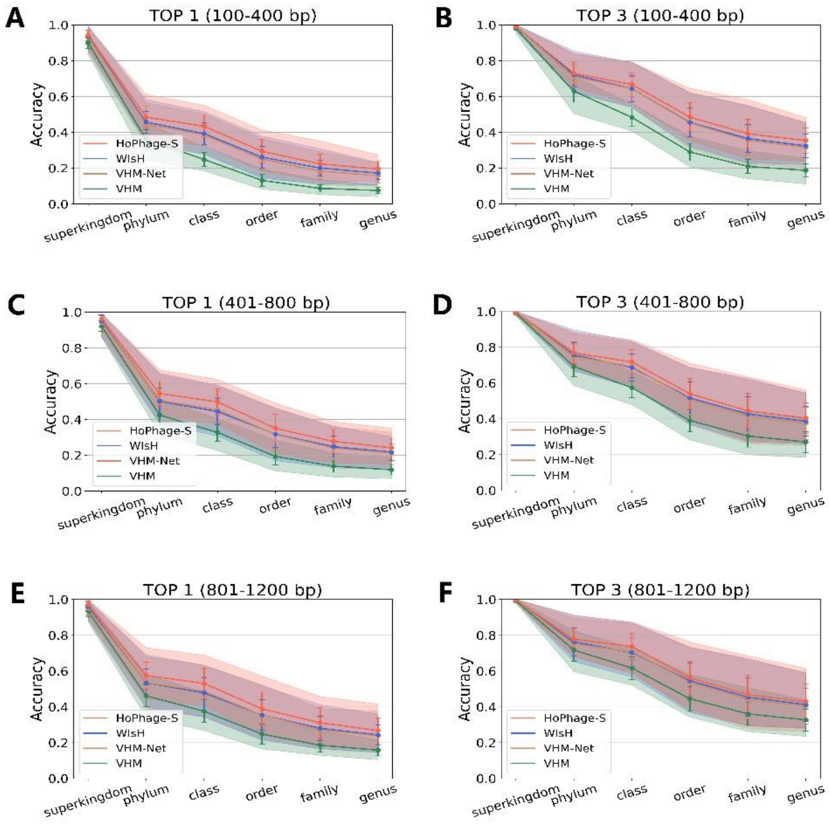


**Fig. S4. Prediction accuracy of HoPhage-S at the different taxonomic levels within the host range of 50 genera and comparison with related tools**. A, C, E are the accuracies of the top 1 prediction of host genus of HoPhage-S, WIsH, VHM-Net and VHM at each taxonomic level on 100-400, 401-800, 801-1200bp, respectively. B, D, F are the results for the top 3 predictions.

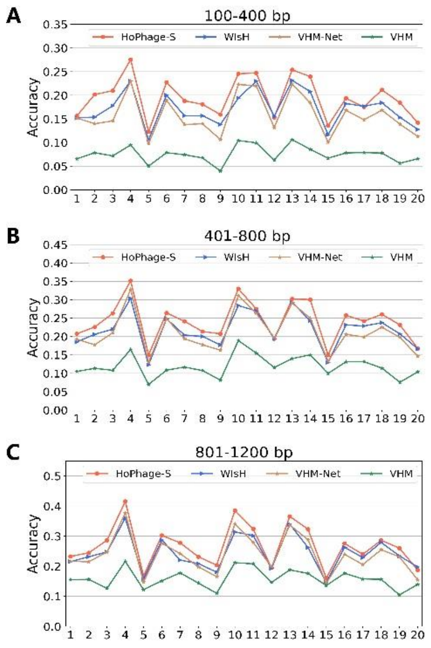


**Fig. S5. Prediction accuracy of HoPhage-S at the genus levels within the host range of 50 genera and comparison with related tools**. A, B, C are the accuracies of the top 1 prediction of host genus of HoPhage-S, WIsH, VHM-Net and VHM at the genus level on 100-400, 401-800, 801-1200bp, respectively. The x-axis represents different random orders.

### 3. Weight selection when integrating HoPhage-G and HoPhage-S

As introduced in the main text, we integrated HoPhage-G and HoPhage-S by calculating the weighted average score of these two modules. As mentioned before, pairs were divided into three categories according to the HoPhage-G score (i.e. Score_Gmax) and subsequently different strategies were used to predict the host. We first counted the distribution of Score_Gmax of phage fragments, the results are shown in Table S2. As the length of the query phage fragments increased, the proportion of high scores also gradually increased, which signified that HoPhage-G can obtain more reliable prediction results for longer sequences.

**Table S2. The distribution of Score_Gmax**.

| The maximum score of HoPhage-G | Score_Gmax ≥ 0.8 | 0.8 > Score_Gmax ≥ 0.4 | 0.4 > Score_Gmax |
|---|---|---|---|
| 100-400 bp | 15.94% | 73.66% | 10.40% |
| 401-800 bp | 32.84% | 66.52% | 0.64% |
| 801-1200 bp | 40.50% | 59.24% | 0.26% |

To determine the best weight of these two modules, we compared the performance of HoPhage with different weights. Table S3 shows the genus accuracies when using different weight strategies to calculate the weighted average score, including uniform weights and stepped weights. In the end, the stepped weighting strategy achieved the best performance, which was conceivable since the level of score represents its reliability, the higher score, the better reliability. However, when the weight of HoPhage-S was between 0.125-0.5, different weights had little effect on the results. Therefore, we recommend that users choose weights according to the actual situation of their data. For example, when the sequence length is long enough, it is better to assign a larger weight to HoPhage-S. Some other suggestions are put forward in the real data application part later. In order to reduce the preference of HoPhage, we then set the weights of both modules to 0.5 to evaluate the model performance and compared it with other tools.

**Table S3. Prediction accuracy of HoPhage at the genus level with different weights**.

| **Weight** | **0** | **0.125** | **0.25** | **0.5** | **0.75** | **0.875** | **1** | **stepped** |
|---|---|---|---|---|---|---|---|---|
| **100-400 bp** | 41.85% | 42.22% | 42.47% | 42.12% | 39.54% | 37.14% | 33.95% | **42.64%** |
| | ±7.15% | ±7.14% | ±6.97% | ±6.56% | ±6.44% | ±6.42% | ±6.09% | **±6.94%** |
| **401-800 bp** | 49.94% | 50.82% | 51.57% | 50.80% | 48.73% | 47.46% | 45.77% | **51.65%** |
| | ±6.53% | ±6.48% | ±6.26% | ±6.05% | ±6.30% | ±6.49% | ±6.49% | **±6.19%** |
| **801-1200 bp** | 55.28% | 55.64% | 56.11% | 55.80% | 54.05% | 52.89% | 51.60% | **56.35%** |
| | ±6.16% | ±6.69% | ±6.67% | ±6.48% | ±6.73% | ±6.92% | ±6.96% | **±6.45%** |

The constant weight is the weight of HoPhage-S, hence the weight of HoPhage-G is (1-weight). The 'stepped' means that the weight used for three categories divided by Score_Gmax is different. S weight becomes larger ss Score_Gmax becomes smaller, which are 0.125, 0.25, and 0.5 respectively.
